# Converging paths: Harnessing ensemble modelling to predict human wild pig conflict risk zones in Tamil Nadu

**DOI:** 10.1101/2023.12.20.572717

**Authors:** Thekke Thumbath Shameer, Priyambada Routray, A Udhayan, Nihar Ranjan, Mannika Govindan Ganesan, Arulmani Manimozhi, Dhayanithi Vasantha Kumari

## Abstract

Growing human populations and human activity intensify human-wildlife conflicts, particularly with wild pigs (*Sus scrofa*), necessitating consideration of both human and wildlife requirements. This complexity demands a comprehensive assessment of causes, impacts, and mitigation strategies for such conflicts. In order to address these issues, we conducted studies across the forest divisions of Tamil Nadu by collecting secondary data on conflicts from 2016 to 2021. Out of the 3301 incidents we collected, 94.4% were specifically related to crop damage and occurred mostly during the month of December, followed by November. Dharmapuri Forest Division was found to be the highest contributor to overall conflicts. Temporal analysis revealed conflict peaks in December, followed by November and September. Using ensemble modelling we predicted a conflict risk zone of approximately 49,223 km2, which represents 37.8% of the total area of Tamil Nadu. Variables like human modification index and mean annual temperature had the highest contribution to model performance. Our model’s projections indicate that areas with cultivated lands in close proximity to the fringes of forests, especially in regions where there is a greater degree of human modification, are associated with heightened levels of conflict risk. This study provides a concise overview of a comprehensive study on human-wild pig conflict, encompassing an exploration of conflict triggers and ecological impacts in the state of Tamil Nadu. The study offers a comprehensive roadmap for managing human-wild pig conflicts in Tamil Nadu, highlighting key triggers, ecological impacts, and conflict risk factors.

## Introduction

Human population growth and subsequent development near natural habitats have raised various negative consequences that have substantially increased human-wildlife conflict (HWC) (Karami and Tavakoli 2023; Nhyus 2016). HWC is a much-discussed term that occurs when the needs of humans and wildlife overlap, resulting in a negative type of interaction for both. (Mekonen, 2020; Dickman, 2010; Naughton-Treves et al. 1998). The escalation of HWC has emerged as a prominent and pressing threat to biodiversity conservation, exhibiting increasing frequency and severity across various regions of the globe (Mekonen, 2020; Karanth et al. 2018; Amaja et al. 2016). HWC can take various forms, such as crop damage, property destruction, and both human and animal casualties. With the increased dependency of humans on natural resources and overexploitation of natural habitats, incidents of HWC are likely to increase, making it a complex issue to address (Rawat et al. 2021; Sharma et al. 2021; Woodroffe, 2005). HWC is driven by multiple factors, including habitat loss and fragmentation, changes in land use, forest cover, agricultural practices, human encroachment into wildlife habitat, and poaching (Sharma et.al. 2020). These factors can result in competition for resources such as food and water. It can lead to direct or indirect interactions between humans and wildlife (Treves 2009). Unfortunately, these interactions have mostly been negative and have caused fatal consequences for humans and wildlife as well. In addition, social and economic factors can also influence HWC, such as overwhelming population growth, poverty, and lack of alternative livelihood options (Nyhus 2016). Understanding the complex drivers of human wildlife conflict is crucial for developing efficacious management strategies that foster coexistence between humans and wildlife while ensuring the welfare of biodiversity and the well-being of local communities (Treves 2014).

Wild pigs come under the generalised species category, which is why they are highly flexible, prolific and distributed over many parts of the world (Lowe 2000). Wild pigs are opportunistic omnivores, feeding on a large selection of foods, starting with roots, tubers, insects, small mammals, and even carrion Schley and Roper (2003). Wild boars also possess a flexible feeding strategy and have a tendency to exploit locally abundant food sources. However, this behavior results in conflicts with humans when they come into contact with agricultural crops or other human-affected areas (Schley et al. 2008). Wild pigs attain reproductive maturation at a very young age and have high reproductive potential and a large litter size (West et al. 2009). These criteria have made wild pigs a significant economic threat in many parts of the world and even considered pests. Human wild pig conflict (HWPC) is an evolving issue that has attracted increasing attention in recent years. Wild pigs are highly versatile and intelligent animals that cause significant damage to crops, property, and ecosystems as well (Massei, 2015). They can transmit diseases to humans and livestock. As human populations keep on expanding, which results in encroaching and fragmenting wild pig habitats, the frequency of human-wild pig conflict is likely to escalate Barrios-Garcia and Ballari (2012). It is evident that HWPC is a key issue in India, particularly in the southern parts of the country, which makes it of utmost importance to address in order to stop putting human lives and properties in danger. The amplification in the population of wild pigs without a proportional growth in forest area has resulted in HWPC, which poses a threat to both subsistence and commercial agriculture. Out of all the different types of conflicts, crop damage and human injury are typical outcomes of HWPC. Several studies have recognized habitat loss and fragmentation, urbanization, and agricultural practices as the fundamental drivers of HWPC (Milda et al. 2023; Senthilkumar et al. 2016; Chauhan et al. 2009).

Species distribution modeling (SDM) is one of the most powerful tools in conservation planning, and researchers have been implementing it in various studies (Shameer et al. 2023; Shameer et al. 2022; Shameer et al. 2021). The ensemble modeling framework suggested by Araujo and New (2007) has gained popularity for predicting reliable SDMs. Ensemble modeling is a powerful approach for improving the accuracy and robustness of species distribution models (SDMs), which involves fitting multiple models using different algorithms, predictors, or modeling techniques and combining the predictions using a weighted average or other aggregation method. This approach can help reduce the effects of overfitting, account for model uncertainty, and capture the complexity of the relationship between species and the environment. The “sdm” package Naimi and Araujo (2016) implemented on the R platform, provides multiple combinations of algorithms for ensemble modeling. This enables users to create a set of candidate models, evaluate their performance, and combine them into an ensemble using various methods such as boosting, bagging, or random forests. Ensemble modeling has been proven to improve the accuracy and reliability of SDMs in many applications, such as predicting the distribution of species, modeling the effects of climate change on biodiversity, identifying priority areas for conservation, and predicting HWC (Karami and Tavakoli, 2022; Ahmed et al. 2021; Mpakairiet al. 2018).

Understanding the regional spatio-temporal patterns and identifying the triggers of HWPC can contribute to the development of effective management strategies, mitigating the impacts on farmers and communities. By gaining deeper insights into the behavior, ecology, and socio-economic factors associated with conflicts, we can establish sustainable and humane solutions to address this pressing issue. These studies will also shed light on potential risks and unintended consequences linked to different management strategies, highlight knowledge loopholes, and promote safe and competent management practices. The outcomes of such research can fill the gaps in the creation of management plans, facilitate stakeholder engagement, garner support for reliable conflict management, and ultimately foster coexistence between humans and wildlife.

Hence, conducting systematic research on HWPC is crucial for developing successful management strategies, evaluating their efficacy, uncovering unanswered questions, and providing evidence-based information to guide the decision-making process and policy development in the right direction. This knowledge can help develop a targeted management approach that can mitigate negative impacts on agriculture, human health and safety, cultural practices, and conservation efforts. Ultimately, long term monitoring is essential to ensure that management decisions are evidence-backed and proficient in resolving HWPC. Accordingly, we conducted a study in Tamil Nadu by collecting data on HWPC that were reported during 2016-2021 from the forest department records and through field surveys to i) understand the distribution of HWPC and their temporal pattern, ii) identify the drivers of HWPC, and iii) predict the conflict risk zones over the entire landscape of Tamil Nadu.

## 2. METHODS

### 2.1. Study area

The state of Tamil Nadu (approximately 11.1271° N lat and 78.6569° E long) is located in the southernmost region of India. The Bay of Bengal surrounds it on the east, the Indian Ocean on the south, Kerala on the west, Karnataka on the northwest, and Andhra Pradesh on the north. The state has varied ranges of terrain, starting from coastal plains to hilly mountain ranges. As mentioned in the National State of Forest Report 2021, Tamil Nadu constitutes a total of 26,451 sq km of forest cover, which is about 17.41% of the total state geographical area. This state witnesses an average rainfall of about 800 mm to 1200 mm, and the annual average temperature ranges from approximately 25°C to 32°C. Since Tamil Nadu experiences various types of climatic conditions that support a diverse range of forest types, starting with tropical wet evergreen forest, tropical semi-evergreen forest, and ending with sub-tropical hill forest and montane wet temperate forest, these many varied forests hold a diverse range of flora and fauna.

### 2.2. Data collection

We collected data from 2016–2021 on HWC incidents across the forest divisions of Tamil Nadu from both secondary sources and field visits. The data had information about HWPC such as date of occurrence, latitude and longitude, conflict species and type of conflict (i.e. crop damage, human injury, and human death), crop name, and compensation details. While visiting the conflict location, we collected information about the frequency, terrain information, and any other ground factors that influence HWC in the region. We engaged in interactions with a few victims of conflicts to gain insights into the victim’s perspective of the incident. Additionally, we discussed with forest officials and farmers to comprehend the mitigation measures employed and their effectiveness.

### 2.3. Data analysis

We entered the data into a spreadsheet. Using the pivot table tool in Excel, we sorted the frequency of conflict by forest division, crop, and type of conflict. We visualised division-wise conflict details to understand their spatial distribution. We conducted a thorough temporal analysis on data spanning a defined timeline, encompassing years and months. For the year-wise conflict analysis, we organized conflict events into tables corresponding to each year and depicted the results through line graphs. This approach allowed us to discern the months characterized by heightened conflict occurrences. To visualize the monthly conflict patterns, we aggregated data on a monthly basis and represented the conflict incidents using bar graphs. We prepared bar charts for visualising the above data using the package ggplot2 (Wickham 2016).

### 2.4. Data thinning

We obtained a total of 3363 independent conflict events from the data mentioned above. To reduce the spatial autocorrelation among the occurrences, we performed a spatial thinning process using the package “spThin” (Aiello-Lammenset al. 2015) in the R platform. Spatial thinning stands as the best method to reduce spatial sampling biases. This strategy involves the selective removal of data while maintaining crucial information in order to mitigate the influence of sampling biases. The thin function in the package spThin employs a randomization approach and returns a maximum number of records within a thinning distance while analysed with sufficient iterations. We removed the conflict records that exhibited spatial autocorrelation and consolidated multiple occurrences. As a result, occurrence records decreased from 3363 to 864 (Fig. 1).

**Fig. 1.**
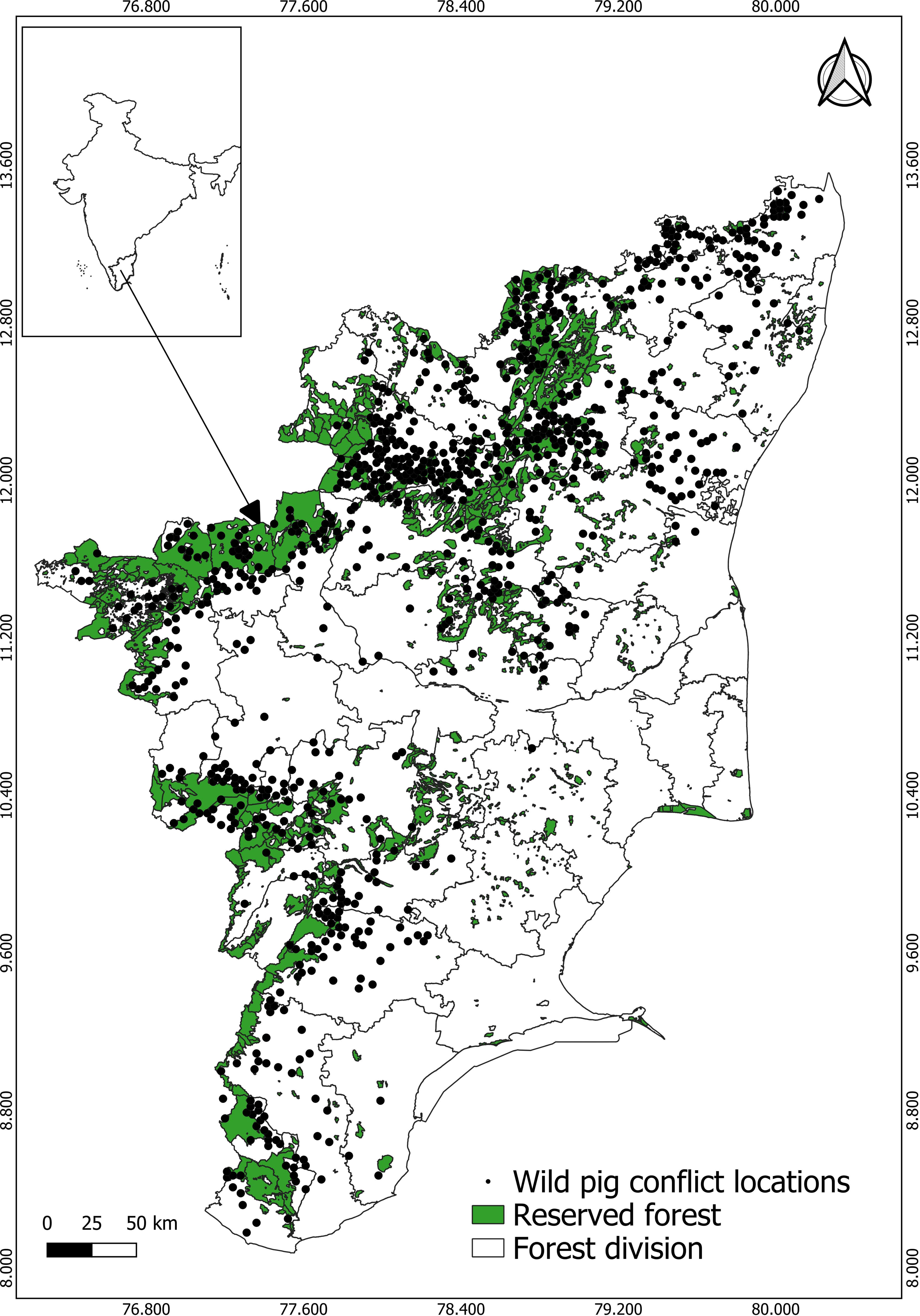
The study area showing the conflict locations is represented in black, while the reserved forest areas are highlighted in green and overlaid on the forest division boundaries of Tamil Nadu, Southern India.

### 2.5. Environmental variables

We choose three bio clime variables such as BIO1 (Annual Mean Temperature), BIO3 (Isothermality), BIO12 (Annual Precipitation) downloaded from Worldclim (http://www.worldclim.org) (Hijmans et al. 2005). We used the SRTM 30m digital elevation model (DEM) by Farr et al. 2007 and calculated the terrain ruggedness index (TRI) using the r package raster and rgdal (Bivand et al. 2023). We further extracted the layers such as forest cover, croplands, build up areas from the ESA World Cover 2022 data base in 30 m resolution using httr (ESA World Cover project / Contains modified Copernicus Sentinel data (2020) processed by the ESA World Cover consortium) and raster (Hijmans, 2012) packages. Further, we converted these rasters to polygons and calculated their Euclidean distances using the package rgeos (Bivand and Rundel, 2021). Normalized Difference Vegetation Index (NDVI) was downloaded from the MODIS (modis.gsfc.nasa.gov) database, and its average was calculated from 2016–2021 of our study period using the Raster package. The water layers were downloaded from Humanitarian OpenStreetMap (https://data.humdata.org/) and their Euclidean distance was calculated. We further downloaded Global Human Modification of Terrestrial Systems (HMI), v1 (2016), which provides a cumulative measure of the human modification of terrestrial lands across the globe at a 1-km resolution (Kennedy et al. 2020). We rescaled all environmental covariates to 1 km resolution using a raster package in the R platform. To address the issue of multicollinearity between variables, we performed Pearson correlation coefficient analysis with a threshold of 0.75 using the package ggplot2 with the function ggcorrplot. We did not find any collinearity between the variables; hence, all the variables were retained and used for the modeling.

### 2.6. Modeling procedure

We generated pseudo absence records equal to our presence records, totalling 864 points, using the gRandom method in the sdm package. We employed nine candidate models available in the sdm package, including Generalized Linear Models (GLM), Boosted Regression Trees (BRT), Random Forests (RF), Flexible Discriminant Analysis (FDA), Maximum Entropy (MaxEnt), Classification and Regression Trees (CART), Multivariate Adaptive Regression Splines (MARS), Support Vector Machines (SVM), and Generalized Additive Models (GAM), to build an ensemble model. Out of the 865 conflict occurrence records, we randomly selected 30% (259 points) for testing the accuracy of the models, while the remaining 70% (606 points) we used for training. We repeated this process five times to calculate the mean values of sensitivity, specificity, True Skill Statistics (TSS), kappa and Area under the Curve (AUC) thereby evaluating the accuracy of the models. To ensure unbiased predictive accuracy with low variance, we employed the bootstrapping replication method, following previous studies (Ahmed et al. 2021; Harrell et al. 1996; Lima et al. 2019), to run the individual algorithms. The independent algorithms were then ensembled using the weighted averaging method, with TSS as the evaluation statistic and a threshold value of maximum sensitivity + sensitivity. We converted the ensemble raster outputs to binary format using the raster package to prepare the predicted conflict risk zone. Finally, we used QGIS 3.30.1 to visualize and generate the final map, combining all of the ensemble model’s outputs and information. The models output have been categorized into four distinct categories to highlight the conflict risk zones. Output values falling between 0 and 0.25 are classified as “Very Low” conflict zones, those between 0.25 and 0.50 as “Low” conflict zones, values between 0.50 and 0.75 as “High” conflict zones, and finally, values ranging from 0.75 to 1.00 are categorized as “Very High” conflict zones. The next step involved calculating the area of conflict risk zones. This was achieved by summing up the total areas designated as “High” and “Very High” conflict zones.

## 3. RESULTS

### 3.1. Data profile of HWPC

According to the data, we retrieved 3331 wild pig conflict incidents. Out of which 3117 (94.4%) crop damage cases, 13 (0.39%) human deaths, 161 (4.9%) human injuries, 4 (0.1%) livestock depredations, and 6 (0.2%) property damage cases. Out of all wild pigs preferred crops such as groundnut (23.0%) and paddy crops (19.9%), while on the other hand, they moderately favoured sugarcane (10.0%), tapioca (7.82%), maize (9.5%), banana (4.6%), and corn (6.0%) (Fig. 5).

### 3.2. Forest Division-wise Distribution of HWPC

Out of 50 forest divisions, 40 have contributed to wild pig conflict incidences (Fig. 2). Out of those 3301 records, the highest number of conflicts were found to be in Dharmapuri division (20.8%), followed by Thiruvallur (16.7%). KMTR Ambasamudram (8.3%), Tiruvanamalai (7.4%), and Tiruppur (5.1%) divisions showed moderate conflicts. Divisions like Trichy (0.6%) and Kanyakumari (0.4%) showed the fewest numbers of conflicts. Other divisions, such as Gudalur (0.2%), Namakkal (0.03%), etc. showed significantly lower numbers of conflicts.

**Fig. 2.**
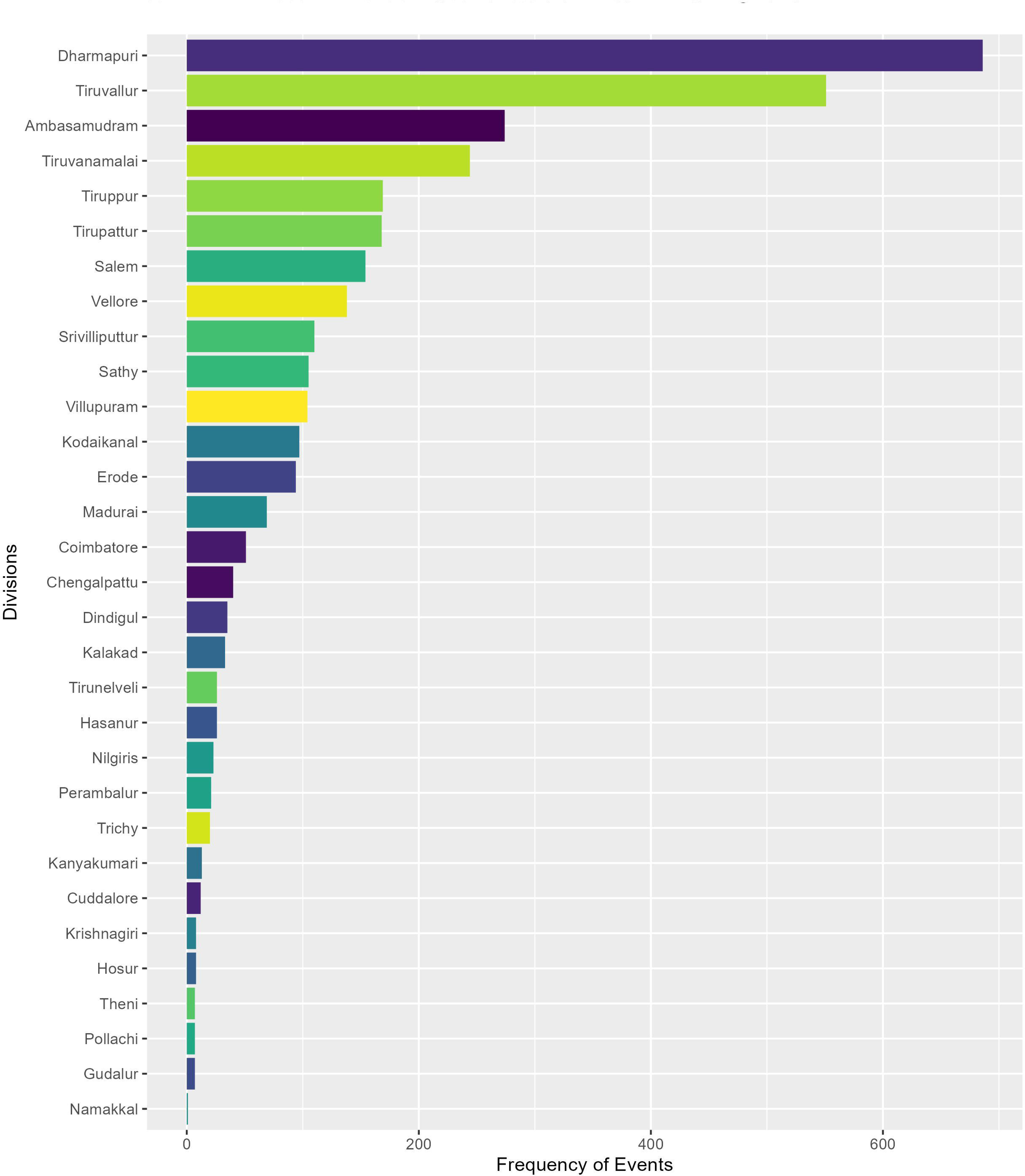
The bar graph illustrates the distribution of conflicts across forest divisions in Tamil Nadu, showcasing the varying levels of HWPC.

### 3.3. Temporal pattern of HWPC

Temporal analysis showed overall, the conflict was high in the month of December, followed by November and September (Fig. 3 b). The January, February, and March month(s) showed moderate amounts of conflict. April had the fewest conflicts, followed by May and June. While comparing the individual year temporal data, we observed the same trends. Our data showed that the year 2020 saw the highest conflict (Fig. 3 a). Crop-wise temporal analysis showed the temporal pattern of each crop preferred by wild pigs (Fig. 4). The banana crop showed that the highest number of conflict incidents occurred from November to March, and July and August also saw moderate amounts of conflict. Mostly they damage Sugarcane crops in the month of January but the trend can start from Sep and last until Mar. Tapioca, on the other hand, can be damaged from June to September. They prefer groundnuts mostly in the month(s) of Aug to Oct, but the conflict occurrence can last from Aug until Mar. Crop damage incidents of corn can start from Oct and last until Jan. Paddy crops showed a pattern of damage from November to March. The maize crop showed a damage pattern from October to January.

**Fig. 3.**
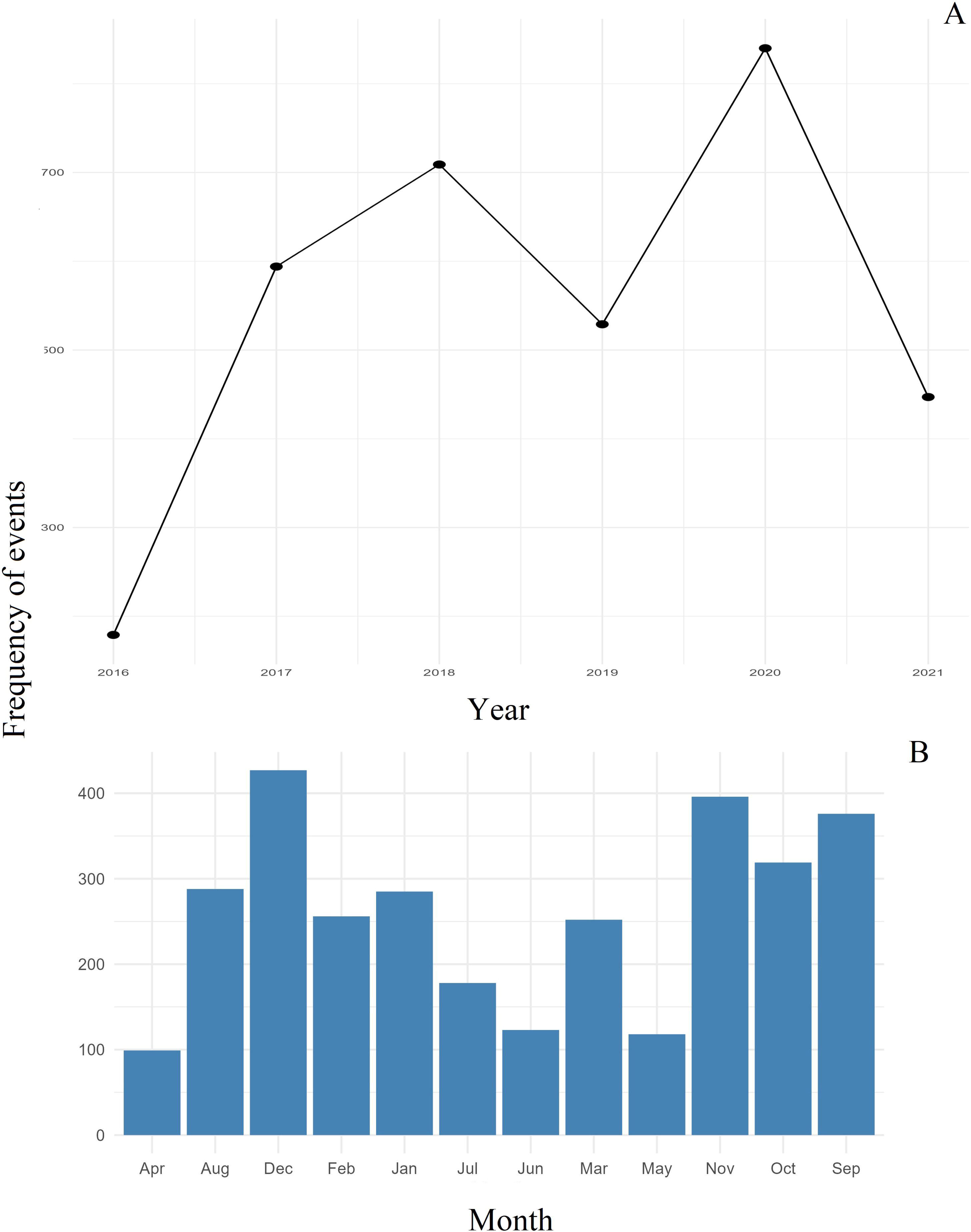
Graph (A) illustrates the year-wise pattern of HWPC, showcasing the fluctuations and trends observed over time. Graph (B) represents the overall temporal pattern of HWPC, providing an overview of the long-term trends and changes in human-wildlife interactions.

**Fig. 4.**
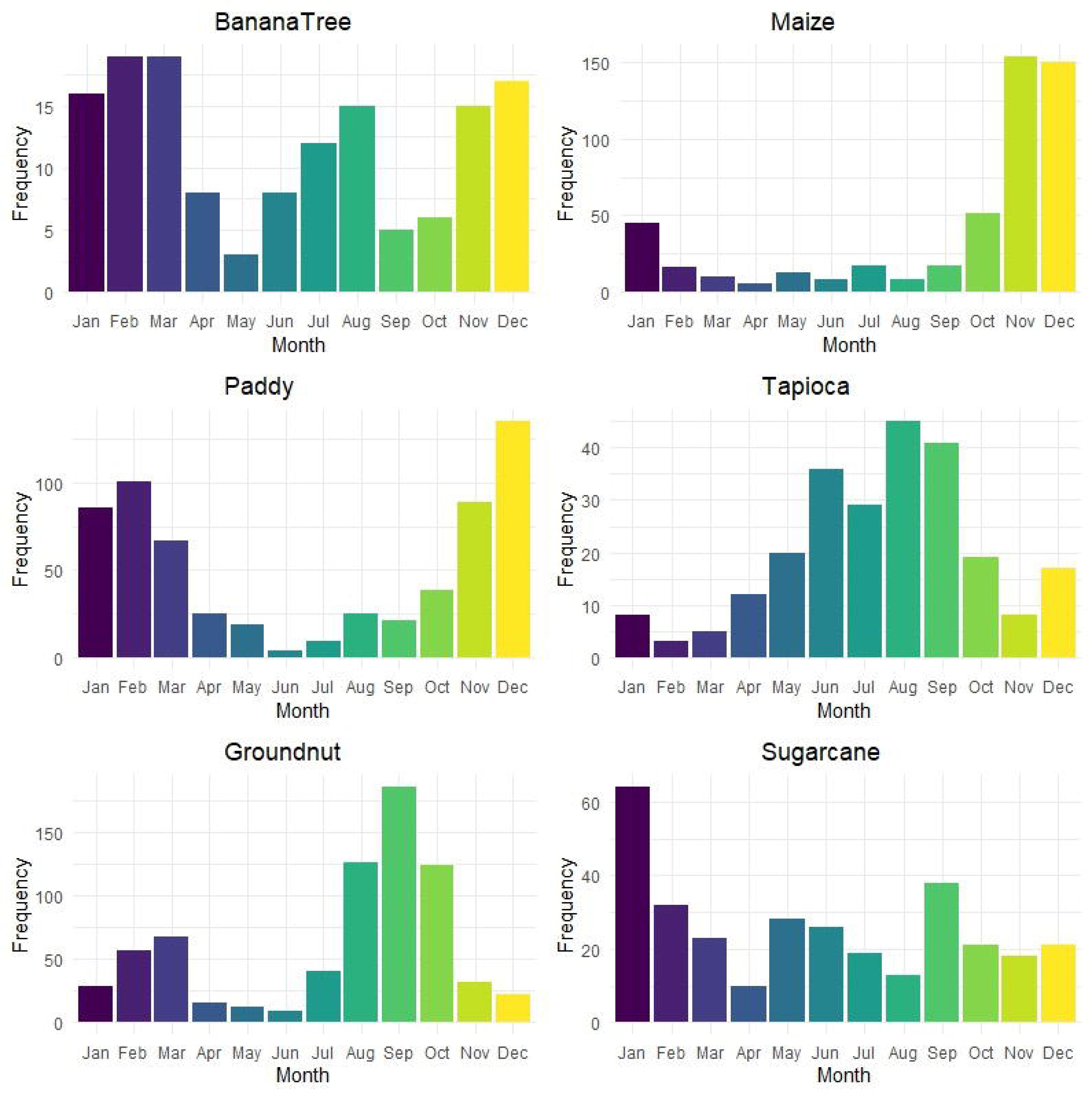
The bar graph displays the temporal pattern of HWPC categorized by different crops, highlighting the variations and trends observed over time.

### 3.4. Model performance and prediction

Supplementary table 1 depicts the performance of models using different evaluation techniques. The top performing model was RF, with an AUC of (0.92), TSS of (0.72) and Kappa of (0.46). The other models, such as SVM, GAM, BRT and MAXENT, MARS also performed well. We also evaluated model accuracy using the receiver operator characteristics (ROC) curve, which has the capacity to show the proportion of the true presence rate and the true absence rate. (Supplementary material 1) shows the ROC curves of all models, which indicate RF as the best model.

Out of the 10-predictor variables, human modification index, mean annual temperature had the highest contribution to model performance (Supplementary material 2). The partial response curve indicated that conflict risk is higher in close proximity to the forest fringes where crop lands and high human modifications occurs (Supplementary material 3). Likewise, we found a higher probability of HWPC occurring with increasing distance towards cropland, built-up, and the HMI. The ensemble model predicts a conflict risk zone covering approximately 49,223 km^2^, which represents 39.67% of the total area of Tamil Nadu. These risk zones were mostly located between the north-east, north-west, and south-western parts of Tamil Nadu (Fig. 6).

**Fig. 5.**
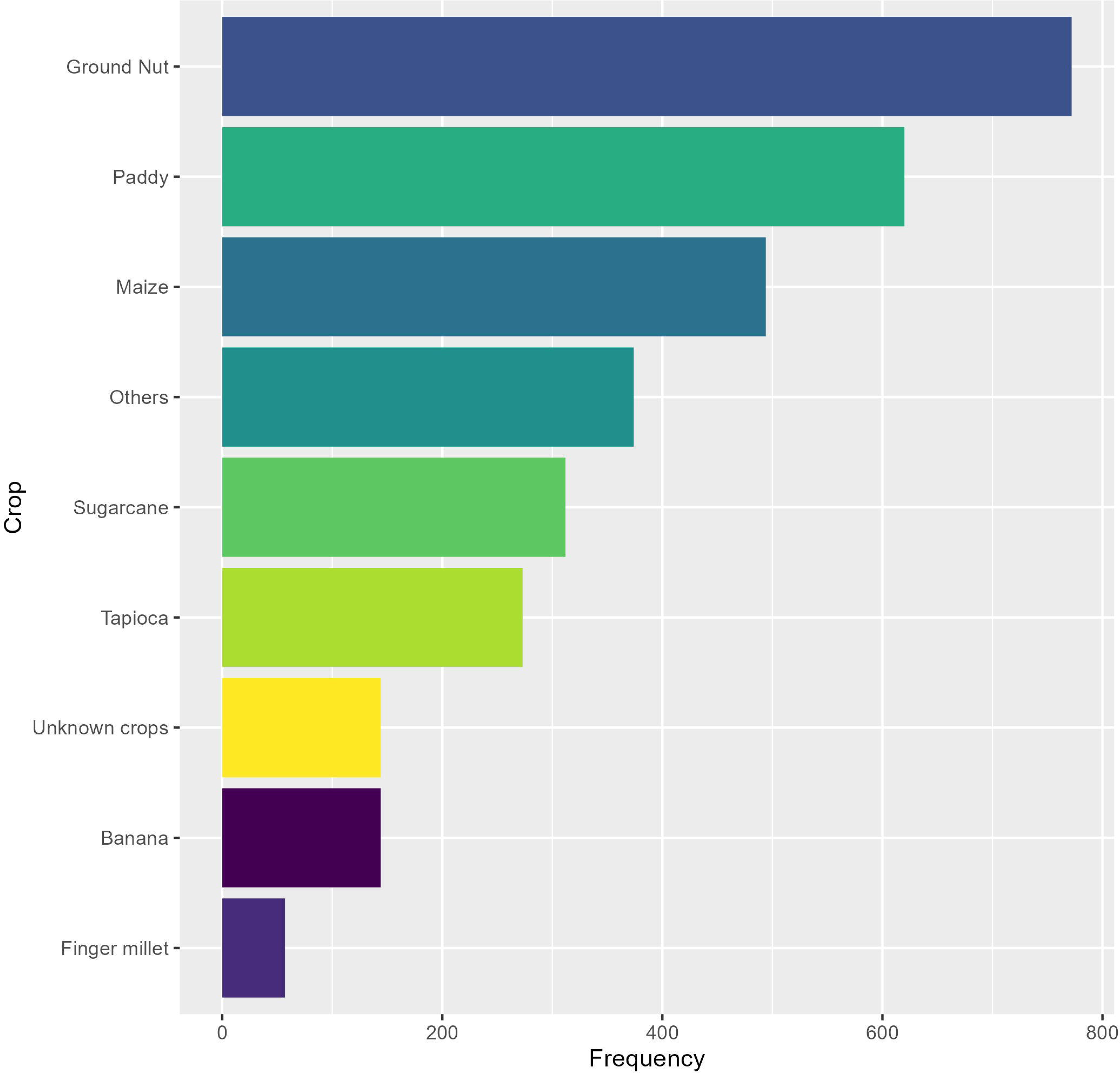
The bar graph illustrates the frequency of HWPC categorized by different crops, providing a visual representation of the occurrence of conflicts associated with each crop.

**Fig. 6.**
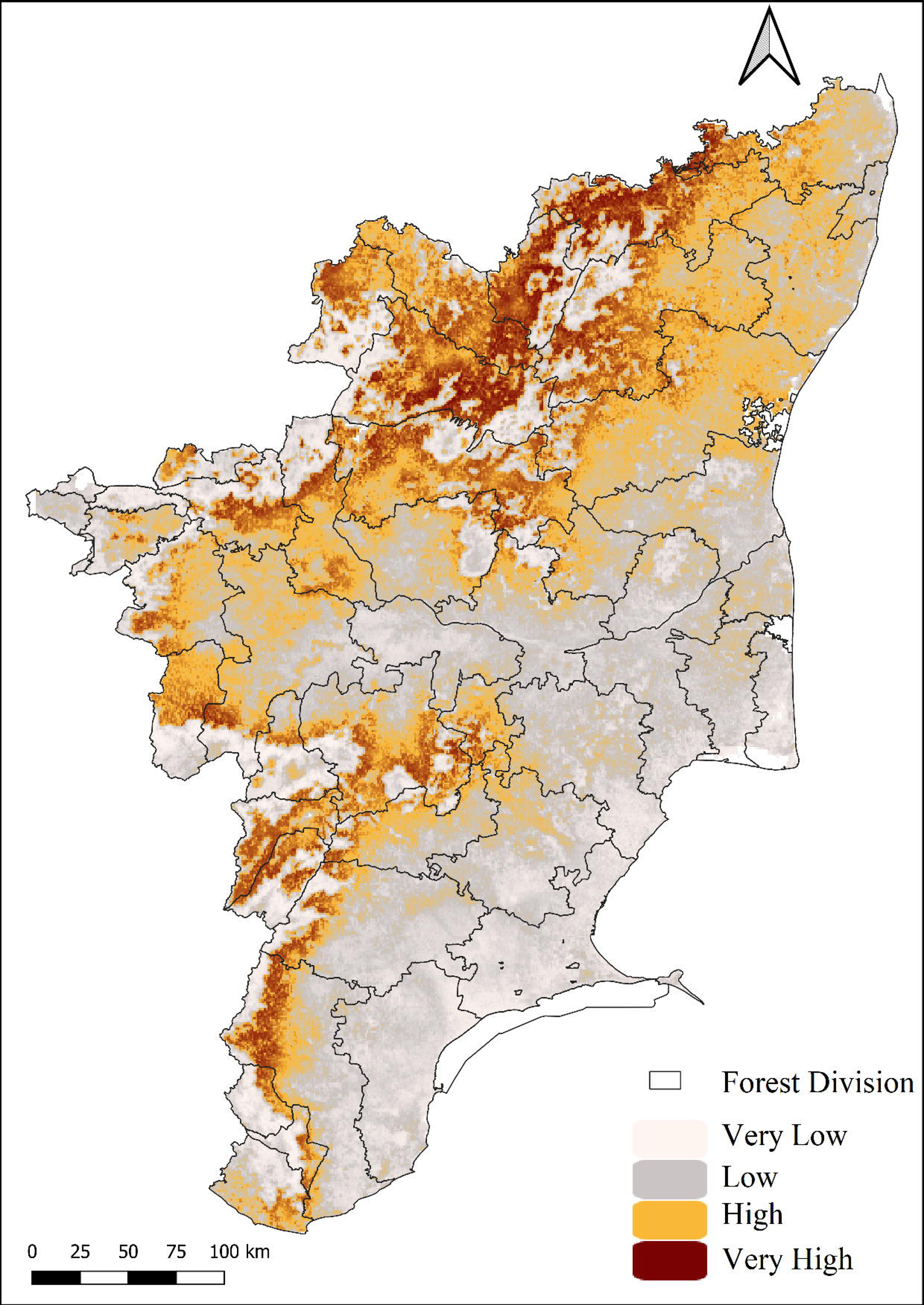
Predicted risk zone of HWPC using ensemble modelling

## 4. DISCUSSION

### 4.1. Extent of crop damage

Based on our results, it becomes evident that crop damage stands out as the most prevalent form of conflict. Crop damage has emerged as a significant global concern in the realm of HWPC (Boyce et al. 2020; McKee et al. 2020; Fischer et al. 2019; Lombardini et al. 2016; Chauhan et al. 2009). This finding aligns with the understanding that wild pigs, being opportunistic omnivores (as noted by Wilcox et al. 2009), have a diverse diet and can consume a wide range of food sources. Although they exhibit a preference for herbs over animal matter, their adaptable nature enables them to exploit various food resources, including crops (Allwin, 2016). This propensity for crop consumption contributes to the significant occurrence of crop damage in the context of HWPC. Wild pigs have experienced a population increase due to the absence of natural predators, coupled with favourable conditions created by human activities near their habitats (Hernandez, 2020; Ickes, 2001; Lewis, 2019; Waithman et al. 1999). These factors provide them with easy access to food and water sources found in human settlements near forest edges (Milda et al. 2023). Consequently, the rise in their population has led to crop damage incidents that can have detrimental effects on the financial conditions of low-income farmers (Pandav et al. 2021). Additionally, human injuries represent the second most common form of conflict (Gulati, 2021; Kabuusu, 2018; Mayer, 2013). Instances of wild pig attacks on humans commonly take place near forest edges, primarily when individuals interact with the animals or when farmers try to deter them from their fields during crop raiding episodes. Wild pigs can exhibit aggression when cornered or perceived as a threat (Chauhan et al. 2009). Although livestock damages are relatively infrequent, occasional instances may occur by mistake. It is important to note that human deaths and property damage are relatively rare, as wild pigs primarily engage in conflicts related to foraging for fodder and do not typically harm humans without provocation. However, there have been isolated cases of human deaths when wild pigs become aggressive and attack individuals they perceive as threats (Mayer, 2013). A consistent pattern emerged in the recorded instances of crop damage, with certain crops that they preferred significantly more. Groundnut (n = 720), paddy (n = 620), sugarcane (n = 312), tapioca (n = 244), maize (n = 297), banana (n = 144), and corn (n = 188) were identified as the primary targets. These crops, commonly cultivated in open fields, are particularly vulnerable to wild pig damage due to their accessibility and the proficiency of wild pigs as diggers. A study by Vasudeva et al. (2015) also confirmed these crops’ susceptibility to damage by wild boars. Additionally, there were approximately 144 types of crop damage by wild pigs reported but not identified.

### 4.2. Distribution Across Forest Divisions and Temporal Trends

We identified that Dharmapuri and Tiruvallur divisions as the major contributors to the overall HWPC records in Tamil Nadu. These divisions carry characteristics like extensive cultivation of cash crops, including groundnut, sugarcane, and paddy, which are highly preferred by wild pigs. Conversely, divisions such as Ambasamudram, Tiruvannamalai, and Tiruppur reported moderate levels of HWPC cases, possibly due to less favourable environmental conditions and fewer suitable croplands compared to Dharmapuri and Tiruvallur divisions. Crop damage is the predominant type of conflict caused by wild pigs, which also showed a significant influence on the temporal pattern of HWPC. We observed that the reported cases of HWPC are most prevalent from the month(s) of August to March, coinciding with the onset of the monsoon season until the end of winter. This aligns with the cultivation seasons of major crops susceptible to wild pig damage as well as their harvesting times. During these month(s), we expect that wild pigs will frequently raid croplands as they seek food resources. An examination of monthly trends in HWPC on a year-by-year basis revealed a consistent pattern. Among the five year(s) of conflict data collected, 2020 stood out with the highest number of reported conflicts. The COVID-19 pandemic could potentially cause this, which enforced restrictions and led to a decrease in human activities. As measures to mitigate conflicts, such as chasing or deterrence, often required personnel strength, and the pandemic halted the operations, wild pigs had greater freedom to roam around human settlements and croplands, resulting in heightened conflicts.

The temporal analysis to understand the seasonal patterns of crop damage revealed interesting trends. For instance, the banana crop exhibited the highest damage occurrence from November to March, which aligns with its typical harvesting time. However, there were notable instances of raids in July and August as well, indicating that young banana plants were also targeted, possibly due to the accessibility of the stem region. Wild pigs are attracted to the edible “root mass” found in young banana trees, making them dig and feed on it. Hence, these factors can explain the seasonal damage patterns of the banana crop. We observed similar patterns for the sugarcane crop, with the highest peak of damage occurring in January. However, wild pigs began raiding sugarcane as early as September and continued until March. Although wild pigs generally prefer mature crops, their raids on young plants may occur due to the availability of food or scarcity elsewhere. Boyce et al. 2020, discovered that the most severe crop damage occurs shortly after planting during the seedling stage and that this can be severe in locations near forests (Mamo et al. 2021). Young crops are an appealing target for foraging and feeding due to their nutritional value and palatability. They are drawn to the tender shoots, leaves, and fruits of young crops, especially if their natural food sources are limited or scarce. This factor may contribute to why wild pigs are damaging young crops in the study area. Groundnut, being a favourite crop for wild pigs, demonstrated a temporal pattern from August to March. These month(s) coincide with the harvesting season for groundnuts. Tapioca, on the other hand, exhibited a slightly different trend, with peak damage cases in August and damages occurring from June to September. As farmers typically harvest tapioca during the monsoon season, it becomes more susceptible to raids during that time. Corn and maize crops displayed a similar temporal pattern, with damage occurring from October to January. These month(s) align with the harvesting season for these crops in the region, making them attractive to wild pigs. Paddy, which we harvest in the winter season, experienced damage mostly from November to March, marking the end of the harvest period. Certain crops have growing seasons that correspond to when wild pig populations are actively foraging. It is important to note that these temporal patterns provide insights into the crop preferences and raiding behaviour of wild pigs. They help in understanding the peak periods of crop vulnerability and can aid in the development of targeted mitigation strategies. Understanding these patterns and factors influencing HWPC can aid in developing effective mitigation strategies.

### 4.3. HWPC risk modelling

Previous studies conducted by Karami and Tavakoli (2023) and Ficetola et al. (2013) have demonstrated the effectiveness of species distribution models (SDMs) in accurately predicting wild pig conflicts. These findings highlight the reliability of SDMs as a robust approach for predicting the risk of HWPC. Therefore, we can consider our modeling approach as a leading method for effectively predicting and assessing the potential risks associated with HWPC. From the ensemble modeling, it is clear that certain variables made a significant contribution to the model’s performance. Of the ten predictor variables analysed, variables such as human modification index, mean annual temperature, digital elevation model and distance to crop land stood out as having the highest contribution and influence in the HWPC model. These variables likely played a crucial role in influencing the occurrence and intensity of HWPC. The partial response curve indicates that the human settlements in proximity to wild pig habitats or potential foraging areas is likely to have increasing trends in HWPC. Several studies indicate that crop damages by wild pigs are usually happening at the forest fringes (Liu et al. 2019; Thurfjell et al. 2009; Jin et al. 2021). Forest fringes provide a favourable habitat for wild pigs due to the combination of forest cover and nearby agricultural fields. We know wild pigs adapt well to such heterogeneous landscapes, utilising both natural and human-altered environments. The contribution of DEM provides information about topographical features that influence wild pig movement patterns and their accessibility to human-dominated areas. The mean annual temperature could affect the distribution and behaviour of wild pigs, potentially influencing their interactions with humans. Studies suggest that there is a strong correlation between temperature and the distribution of wild pigs. McClure et al. 2015 noted that the distribution of wild pigs was most strongly limited by cold temperatures, and a high probability of occurrence was associated with frequent high temperatures, up to a high threshold. They also reported that they are also likely to occur where potential home ranges have higher habitat heterogeneity, providing access to multiple key resources including water, forage, and cover. A strong correlation is evident between elevated NDVI values and an higher likelihood of conflict events. Considering that NDVI serves as an indicator of vegetation density and health within specific geographic areas, this correlation underscores the pivotal role of substantial vegetation presence within conflict-prone zones. In conflict zones, croplands with abundant vegetation cover, such as dense crops or areas with tall grasses, can potentially serve as favourable refuge habitats for wild pigs, offering protection from predators and human disturbances (Barasona et al. 2021). Similarly, a higher likelihood of conflict risk was observed to correlate with an increasing human modification index. Rutten et al. (2019) revealed that wild pigs demonstrate remarkable behavioural and physiological adaptability in response to human-dominated landscapes. This suggests that the proximity to these features or land uses may increase the likelihood of conflicts between humans and wildlife, potentially due to factors such as resource availability, human disturbance, or the attraction of wildlife to these areas, which is also evident from several HWC studies (Sharma et al. 2020; Huang et al. 2018; Markovchick-Nicholls et al. 2007). These findings highlight the importance of considering landscape features and land use patterns when understanding and managing HWPC. They suggest that conserving and managing forested areas while also considering the spatial relationships between human settlements, croplands, build-up areas, and wildlife habitats can be crucial for mitigating conflicts and promoting coexistence between humans and wildlife.

### 4.4. Management implications

The management of HWPC has prompted a range of strategies employed by both the forest department and local farmers. The existing measures encompass cloth fencing, low-height electric fencing, and chasing, compensation, and awareness campaigns. Notably, among these, cloth fencing has demonstrated a degree of effectiveness, as reported by forest officials and farmers. Additionally, certain areas have experimented with Bio repellent oil as a mitigation measure, although its utility is limited due to its high cost and short duration of effectiveness. Addressing the complexity of the issue demands a multi-faceted approach, drawing on insights from various disciplines. This entails active collaboration between local farmers, panchayats, police departments, and medical departments to expedite compensation processes. A strategic proposition involves the introduction of insurance options for farmers or the provision of subsidies to those unable to install protective fences. Emphasizing community engagement is crucial, given the interconnected nature of the challenge; securing one farm might inadvertently lead to wild pig raids on neighbouring crop fields, thereby escalating wild pig conflicts. To address this, a comprehensive solution is proposed: the establishment of fencing encompassing entire community croplands, facilitated by NGOs or government initiatives. In this arrangement, farmers would assume responsibility for fence maintenance, adapting it as necessary. By fostering this collaborative model, a more sustainable approach to managing HWPC can be achieved, minimizing conflicts while promoting harmonious coexistence between humans and wildlife.

During our discussions with forest officials, it became evident that a more streamlined approach is needed for the allocation of funds. The current allocation doesn’t adequately cater to the varying degrees of challenges faced by different ranges. It’s imperative to adopt a more nuanced funding strategy that takes these variations into account. To enhance the efficiency of wildlife conflict management, forest officials have highlighted the necessity of revising the allocated budget. This is particularly critical for ranges that engage in frequent nightly patrolling and pursuit activities. A recurring issue arises from the promotion of watchers to guard positions, resulting in vacant watcher and anti-poaching watcher (APW) roles. To address this, a consistent recruitment process for these positions needs to be established. Furthermore, there’s a disparity in resources and staffing between larger and smaller ranges. Some ranges lack sufficient vehicles and fuel, impacting their operational capabilities. It’s worth exploring the allocation of additional resources to address these discrepancies. Additionally, for larger ranges, there’s potential to optimize management by dividing them into smaller units and recruiting new staff accordingly. This approach could significantly enhance conflict resolution efficiency. In pursuit of a comprehensive solution, there’s a suggestion to empower and educate local tribes. This can be achieved through training programs that enable them to form committees dedicated to conservation and sustainability. By involving the local communities, we not only disseminate crucial knowledge about wildlife conservation but also create an avenue for passing on this awareness to the younger generation. This approach has the potential to cultivate a generation of wildlife guardians who understand and value the significance of conservation efforts.

Engaging with farmers, we have unearthed certain limitations in the current compensation structure, which appears to be somewhat superficial. For instance, the prevailing practice of offering a flat compensation of 500/- per coconut tree, regardless of its maturity, is being questioned. Farmers propose a more context-sensitive approach by factoring in the developmental stage of the crop when determining compensation. This suggestion applies not only to coconut trees but extends to other crops as well. Implementing such an approach could render compensation more reflective of the actual losses suffered by farmers.

Drawing inspiration from global and national examples, an innovative measure involves using human scalp hair (HSH) as a deterrent. This method has demonstrated effectiveness in different Indian states. The hair, when ingested by wild pigs, causes irritation and disrupts their activities. Pigs encountering individuals with hair in their snouts produce alarm calls, deterring others from venturing into crop lands (Rao et al. 2015). Likewise, other practices from various regions have been identified, such as deploying solutions derived from wild pig dung and burning dried dung cakes to minimize territorial conflicts (Rao et al. 2015).

It’s recommended that mitigation strategies be tailored to the specific location, household socioeconomic status, wildlife species, and regional policies (Karanth et al. 2018). Numerous researchers have put forth additional suggestions, including the use of wire mesh fencing (Thapa, 2010), pig-proof barriers (Chauhan et al. 2009), and cultivating less preferred crops or altering cropping patterns (Barwal, 2013). Population control strategies have also been explored, as demonstrated by studies (Colomer et al. 2021).

The concept of the “Reserve effect” proposes occasional hunting to disrupt wild pig social organization. Studies suggest that such hunting can potentially mitigate crop damage caused by wild pigs Geisser and Reyer (2004). By leveraging these insights and practices, a more comprehensive and effective approach to managing human-wildlife conflict can be developed, ultimately benefiting both farmers and wildlife conservation efforts.

## Supporting information

The graph presents the Receiver Operating Characteristic (ROC) curve, depicting the performance of the model.

The graph indicates the relative variable importance for correlation and AUC metrics.

The graph shows the Partial Response Curve, indicating the relationship between the independent variables and their impact on the prediction outcome.

## Acknowledgments

The authors acknowledge the Tamil Nadu Forest Department, the PCCF and Chief Wildlife Warden for granting the necessary permissions and all the forest department officials for providing full cooperation and support during our field visits.

## Author Contribution

TTS designed the study. TTS and PR did the fieldwork and analysed the data. TTS and PR constructed figures. TTS and PR wrote the manuscript. U, NR, MGG, AM and DVK mobilized the funds, procured permission, and provided field and logistical support. All authors contributed to the writing of the manuscript.

## Data availability

All the data used in the manuscript is available in the form of graphs or tables. Further inquiries can be directed to the corresponding author.

**Supplementary material 1.** The graph presents the Receiver Operating Characteristic (ROC) curve, depicting the performance of the model.

**Supplementary material 2.** The graph indicates the relative variable importance for correlation and AUC metrics.

**Supplementary material 3.** The graph shows the Partial Response Curve, indicating the relationship between the independent variables and their impact on the prediction outcome.

**Table 1.** displays the evaluation of SDMs utilizing various statistical parameters.

## References

1. Ahmed N, Atzberger C, Zewdie W (2021) Species Distribution Modeling performance and its implication for Sentinel-2-based prediction of invasive Prosopis juliflora in lower Awash River basin, Ethiopia. Ecological Processes, 10(1) 10.1186/s13717-021-00285-6

2. Aiello-Lammens ME, Boria RA, Radosavljevic A, Vilela B, Anderson RP (2015) spThin: an R package for spatial thinning of species occurrence records for use in ecological niche models. Ecography, 38(5), 541–545. 10.1111/ecog.01132

3. Amaja LG, Debela HF, Tariku MG (2016) Assessment of types of damage and causes of human-wildlife conflict in Gera district, south western Ethiopia. Journal of Ecology and The Natural Environment 8(5):49–54. 10.5897/jene2015.0543

4. Araujo M, New M (2007) Ensemble forecasting of species distributions. Trends in Ecology & Evolution, 22(1), 42–47. 10.1016/j.tree.2006.09.010

5. Baral K, Sharma H P, Kunwar R, Morley C, Aryal A, Rimal B, Ji W (2021) Human Wildlife Conflict and Impacts on Livelihood: A Study in Community Forestry System in Mid–Hills of Nepal. Sustainability, 13(23), 13170.

6. Barasona JA, Carpio A, Boadella M, Gortazar C, Piñeiro X, Zumalacárregui C, Viñuela J (2021) Expansion of native wild boar populations is a new threat for semi-arid wetland areas. Ecological Indicators, 125, 107563. 10.1016/j.ecolind.2021.107563

7. Barrios-Garcia MN, Ballari SA (2012) Impact of wild boar (Sus scrofa) in its introduced and native range: a review. Biological Invasions, 14(11), 2283–2300. 10.1007/s10530-012-0229-6

8. Bivand R, Keitt T, Rowlingson B (2023) rgdal: Bindings for the ‘Geospatial’ Data Abstraction Library. Retrieved from http://rgdal.r-forge.r-project.org, https://gdal.org, https://proj.org, https://r-forge.r-project.org/projects/rgdal/

9. Bivand R, Rundel C (2021) rgeos: Interface to Geometry Engine - Open Source (‘GEOS’) Retrieved from https://r-forge.r-project.org/projects/rgeos/, https://libgeos.org, http://rgeos.r-forge.r-project.org/index.html

10. Boyce C, VerCauteren K, Beasley J (2020) Timing and extent of crop damage by wild pigs (Sus scrofa Linnaeus) to corn and peanut fields. Crop Protection, 133, 105131. 10.1016/j.cropro.2020.105131

11. Chauhan NS, Barwal KS, Kumar D, Náhlik A (2009) Human-wild pig conflict in selected states in India and mitigation strategies. Acta Silvatica & Lignaria Hungarica, 5, 189–197.

12. Evans MJ, Hawley JE, Rego PW, Rittenhouse TA. (2014) Exurban land use facilitates human-black bear conflicts. The Journal of Wildlife Management, 78(8), 1477–1485. 10.1002/jwmg.796

13. Colomer J, Rosell C, Rodriguez-Teijeiro JD, Massei G (2021) ‘Reserve effect’: An opportunity to mitigate human-wild boar conflicts. Science of The Total Environment, 799, 148721.

14. Dickman AJ (2010) Complexities of conflict: the importance of considering social factors for effectively resolving human-wildlife conflict. Animal Conservation, 13(5), 458–466. 10.1111/j.1469-1795.2010.00368.x

15. Farr TG, Rosen PA, Caro E, Crippen R, Duren R, Hensley S, Kobrick M, Paller M, Rodriguez E, Roth L, Seal D, Shaffer S, Shimada J, Umland J, Werner M, Oskin M, Burbank D, Alsdorf D (2007) The Shuttle Radar Topography Mission. Reviews of Geophysics, 45(2) 10.1029/2005RG000183

16. Ficetola GF, Bonardi A, Mairota P, Leronni V, Padoa-Schioppa E (2014) Predicting wild boar damages to croplands in a mosaic of agricultural and natural areas. Current Zoology, 60(2), 170–179. 10.1093/czoolo/60.2.170

17. Fischer JW, Greiner K, Lutman MW, Webber BL, Vercauteren KC (2019) Use of unmanned aircraft systems (UAS) and multispectral imagery for quantifying agricultural areas damaged by wild pigs. Crop Protection, 125, 104865. 10.1016/j.cropro.2019.104865

18. Geisser H, Reyer h-u (2004) Efficacy of hunting, feeding, and fencing to reduce crop damage by wild boars. Journal of wildlife management 68(4):939–946

19. Grenouillet G, Buisson L, Casajus N, Lek S (2011) Ensemble modelling of species distribution: the effects of geographical and environmental ranges. Ecography, 34(1), 9–17. 10.1111/j.1600-0587.2010.06152.x

20. Gulati S, Karanth KK, Le NA, Noack F (2021) Human casualties are the dominant cost of human–wildlife conflict in India. Proceedings of the National Academy of Sciences, 118(8) 10.1073/pnas.1921338118

21. Harrell FE, Lee KL, Mark DB. (1996) Multivariable prognostic models: issues in developing models, evaluating assumptions and adequacy, and measuring and reducing errors. Statistics in Medicine, 15(4), 361–387. 10.1002/(SICI)1097-0258(19960229)15:4%3C361::AID-SIM168%3E3.0.CO;2-4

22. Hernández FA, Carr AN, Milleson MP, Merrill HR, Avery ML, Parker BM, Wisely SM (2020) Dispersal and Land Cover Contribute to Pseudorabies Virus Exposure in Invasive Wild Pigs. EcoHealth, 17(4), 498–511. 10.1007/s10393-020-01508-6

23. Hijmans RJ, Cameron SE, Parra JL, Jones PG, Jarvis A (2005) Very high resolution interpolated climate surfaces for global land areas. International Journal of Climatology, 25(15), 1965–1978. 10.1002/joc.1276

24. Hijmans RJ, Etten JV (2012) raster: Geographic analysis and modeling with raster data. R package version 2.0–12. http://CRAN.R-project.org/package=raster

25. Hone J, Atkinson B. (1983) Evaluation of Fencing to Control Feral Pig Movement. Wildlife Research, 10(3), 499. 10.1071/wr9830499

26. Huang C, Li X, Shi L, Jiang X (2018) Patterns of human-wildlife conflict and compensation practices around Daxueshan Nature Reserve, China. Zoological Research, 39(6), 406–412. 10.24272/j.issn.2095-8137.2018.056

27. Ickes K (2001) Hyper-abundance of Native Wild Pigs (Sus scrofa) in a Lowland Dipterocarp Rain Forest of Peninsular Malaysia1. Biotropica, 33(4), 682–690. 10.1111/j.1744-7429.2001.tb00225.x

28. Jin Y, Kong W, Yan H, Bao G, Liu T, Ma Q, Zhang M (2021) Multi-Scale Spatial Prediction of Wild Boar Damage Risk in Hunchun: A Key Tiger Range in China. Animals, 11(4), 1012. 10.3390/ani11041012

29. Kabuusu RM, Amuno JB, Maseruka Y, Macpherson C (2018) Incidence and Risk Factors of Wildlife-Associated Human Injuries in Queen Elizabeth National Park, Uganda: A Retrospective Cohort Study. Journal of Zoonotic Diseases and Public Health, 2(1), 1.

30. Karami P, Tavakoli S (2022) Identification and analysis of areas prone to conflict with wild boar (Sus scrofa) in the vineyards of Malayer County, western Iran. Ecological Modelling, 471, 110039. 10.1016/j.ecolmodel.2022.110039

31. Karanth KK, Gupta S, Vanamamalai A (2018) Compensation payments, procedures and policies towards human-wildlife conflict management: Insights from India. Biological Conservation 227:383–389. 10.1016/j.biocon.2018.07.006

32. Kassambara A (2020) ggcorrplot: Visualization of a Correlation Matrix using ‘ggplot2’. R package version 0.1.3. Retrieved from https://CRAN.R-project.org/package=ggcorrplot

33. Kennedy CM, Oakleaf JR, Theobald DM, Baruch-Mordo S, Kiesecker J (2020) Global Human Modification of Terrestrial Systems. Palisades, New York: NASA Socioeconomic Data and Applications Center (SEDAC) 10.7927/edbc-3z60. Accessed 26/04/2023.

34. Lewis JS, Corn JL, Mayer JJ, Jordan TR, Farnsworth ML, Burdett CL, Miller R S (2019) Historical, current, and potential population size estimates of invasive wild pigs (Sus scrofa) in the United States. Biological Invasions, 21(7), 2373–2384. 10.1007/s10530-019-01983-1

35. Lima TA, Beuchle R, Langner A, Grecchi RC, Griess VC, Achard F (2019) Comparing Sentinel-2 MSI and Landsat 8 OLI Imagery for Monitoring Selective Logging in the Brazilian Amazon. Remote Sensing, 11(8), 961. 10.3390/rs11080961

36. Liu Q, Yan K, Lu YF, Li M, Yan YY (2019) Conflict between wild boars (Sus scrofa) and farmers: distribution, impacts, and suggestions for management of wild boars in the Three Gorges Reservoir Area. Journal of Mountain Science, 16(10), 2404–2416. 10.1007/s11629-019-5453-4

37. Lombardini M, Meriggi A, Fozzi A (2016) Factors influencing wild boar damage to agricultural crops in Sardinia (Italy) Current Zoology, zow099. 10.1093/cz/zow099

38. Lowe S, Browne M, Boudjelas S, De Poorter M (2000) 100 of the world’s worst invasive alien species: a selection from the global invasive species database. The Invasive Species Specialist Group (ISSG), Auckland, 12, 12–14.

39. Mamo A, Lemessa D, Diriba OH, Hunde D (2021) Pattern of crop raiding by wild large mammals and the resultant impacts vary with distances from forests in Southwest Ethiopia. Ecology and Evolution, 11(7), 3203–3209. 10.1002/ece3.7268

40. Markovchick-nicholls L, Regan HM, Deutschman DH, Widyanata A, Martin B, Noreke L, Ann Hunt T (2007) Relationships between Human Disturbance and Wildlife Land Use in Urban Habitat Fragments. Conservation Biology, 22(1), 99–109. 10.1111/j.1523-1739.2007.00846.x

41. Massei G, Kindberg J, Licoppe A, Gačić D, Šprem N, Kamler J, Náhlik A (2015) Wild boar populations up, numbers of hunters down? A review of trends and implications for Europe. Pest Management Science, 71(4), 492–500. 10.1002/ps.3965

42. Mayer JJ (2013) Wildlife Damage Management Conferences-Proceedings (Vol. 151) Retrieved from http://digitalcommons.unl.edu/icwdm_wdmconfproc/151

43. McClure ML, Burdett CL, Farnsworth ML, Lutman MW, Theobald DM, Riggs PD, Miller RS (2015) Modeling and Mapping the Probability of Occurrence of Invasive Wild Pigs across the Contiguous United States. PLOS ONE, 10(8), e0133771. 10.1371/journal.pone.0133771

44. McKee S, Anderson A, Carlisle K, Shwiff SA (2020) Economic estimates of invasive wild pig damage to crops in 12 US states. Crop Protection, 132, 105105. 10.1016/j.cropro.2020.105105

45. Mekonen S (2020) Coexistence between human and wildlife: the nature, causes and mitigations of human wildlife conflict around Bale Mountains National Park, Southeast Ethiopia. BMC Ecology, 20(1). 10.1186/s12898-020-00319-1

46. Milda D, Ramesh T, Kalle R, Gayathri V, Thanikodi M, Ashish K (2022) Factors driving human–wild pig interactions: implications for wildlife conflict management in southern parts of India. Biological Invasions, 25(1), 221–235. 10.1007/s10530-022-02911-6

47. Mpakairi K, Ndaimani H, Vingi K, Madiri TH, Nekatambe T (2018) Ensemble modelling predicts Human Carnivore Conflict for a community adjacent to a protected area in Zimbabwe. African Journal of Ecology, 56(4), 957–963. 10.1111/aje.12526

48. Naimi B, Araújo MB, (2016) sdm: a reproducible and extensible R platform for species distribution modelling. Ecography, 39(4), 368–375. 10.1111/ecog.01881

49. Naughton-Treves L, Treves A, Chapman C, Wrangham R (1998) Temporal patterns of crop-raiding by primates: linking food availability in croplands and adjacent forest. Journal of Applied Ecology, 35(4), 596–606. 10.1046/j.1365-2664.1998.3540596.x

50. Nyhus PJ (2016) Human–Wildlife Conflict and Coexistence. Annual Review of Environment and Resources, 41(1), 143–171. 10.1146/annurev-environ-110615-085634

51. Pandav B, Lakshminarayanan N, Kumar A, Desai A, Lyngkhoi B (2021) Household perceptions and patterns of crop loss by wild pigs in north India. Human-Wildlife Interactions, 15, 1–10.

52. Rao VV, Naresh B, Reddy VR., Sudhakar C, Venkateswarlu P, Rao D R (2015) Traditional management methods used to minimize wild boar (Sus scrofa) damage in different agricultural crops at Telangana state, India. International Journal of Multidisciplinary Research and Development, 2(2), 32–36.

53. Rawat PK, Pant DB, Pant DKK, Pant DP (2021) Geospatial Analysis of Alarmingly Increasing Human-Wildlife Conflict in Jim Corbett National Park’s Ramnagar Urban Buffer Zone: Ecological and Socioeconomic Perspectives. SSRN Electronic Journal. 10.2139/ssrn.3938066

54. Rawat PK, Pant DrB, Pant DrKK, Pant DrP (2021) Geospatial Analysis of Alarmingly Increasing Human-Wildlife Conflict in Jim Corbett National Park’s Ramnagar Urban Buffer Zone: Ecological and Socioeconomic Perspectives. SSRN Electronic Journal. 10.2139/ssrn.3938066

55. Reidy MM, Campbell TA, Hewitt DG (2008) Evaluation of Electric Fencing to Inhibit Feral Pig Movements. Journal of Wildlife Management, 72(4), 1012–1018. 10.2193/2007-158

56. Rutten A, Casaer J, Swinnen KR, Herremans M, Leirs H (2019) Future distribution of wild boar in a highly anthropogenic landscape: Models combining hunting bag and citizen science data. Ecological Modelling, 411, 108804. 10.1016/j.ecolmodel.2019.108804

57. Schley L, Roper TJ (2003) Diet of wild boar Sus scrofa in Western Europe, with particular reference to consumption of agricultural crops. Mammal Review, 33(1), 43–56. 10.1046/j.1365-2907.2003.00010.x

58. Shameer TT, Backer SJ, Nandhini S, Raman S, Mujawar AN, Yogesh J, Sanil, R (2022) How do the sympatric forest mongooses coexist in the Western Ghats landscape? Insights from spatio-temporal approach. Community Ecology, 23(2), 231–245. 10.1007/s42974-022-00101-x

59. Shameer TT, Backer SJ, Yogesh J, Mujawar AN, Ali SZ, Raman S, Sanil R (2021) Phenotypic variations, habitat suitability, and diel activity of the endemic brown palm civets. Geology, Ecology, and Landscapes, 1–12. 10.1080/24749508.2021.1971411

60. Shameer TT, Mungi NA, Backer SJ, Raman S, Reddy SR, Easa PS, Sanil, R (2023) Distribution and conservation status of the endemic Nilgiri marten (Martesgwatkinsii) Mammalia, 0(0) 10.1515/mammalia-2021-0113

61. Sharma P, Chettri N, Uddin K, Wangchuk K, Joshi R, Tandin T, Sharma E (2020) Mapping humanLwildlife conflict hotspots in a transboundary landscape, Eastern Himalaya. Global Ecology and Conservation, 24, e01284.10.1016/j.gecco.2020.e01284

62. Sharma P, Chettri N, Wangchuk K (2021) Human–wildlife conflict in the roof of the world: Understanding multidimensional perspectives through a systematic review. Ecology and Evolution 11(17):11569–11586. 10.1002/ece3.7980

63. Sharma P, Chettri N, Wangchuk K (2021) Human–wildlife conflict in the roof of the world: Understanding multidimensional perspectives through a systematic review. 10.22541/au.162495471.10485884/v1

64. Sherchan R, Bhandari A (2017) Status and trends of human-wildlife conflict: A case study of Lelep and Yamphudin region, Kanchenjunga Conservation Area, Taplejung, Nepal. Conservation Science, 5(1), 19– 25.10.3126/cs.v5i1.24296

65. Thapa S (2010) Effectiveness of crop protection methods against wildlife damage: A case study of two villages at Bardia National Park, Nepal. Crop Protection, 29(11), 1297–1304. 10.1016/j.cropro.2010.06.015

66. Thurfjell H, Spong G, Olsson M, Ericsson G (2015) Avoidance of high traffic levels results in lower risk of wild boar-vehicle accidents. Landscape and Urban Planning, 133, 98–104. 10.1016/j.landurbplan.2014.09.015

67. Treves A, Wallace R B, White S (2009) Participatory Planning of Interventions to Mitigate Humanâ Wildlife Conflicts. Conservation Biology, 23(6), 1577–1587. 10.1111/j.1523-1739.2009.01242.x

68. Treves, A, &Bruskotter, J. (2014) Tolerance for Predatory Wildlife. Science, 344(6183), 476–477. 10.1126/science.1252690

69. Waithman J D, Sweitzer RA, Vuren DV, Drew JD, Brinkhaus AJ, Gardner IA, Boyce WM (1999) Range Expansion, Population Sizes, and Management of Wild Pigs in California. The Journal of Wildlife Management, 63(1), 298. 10.2307/3802513

70. West BC, Cooper AL, Armstrong JB (2009) Managing wild pigs: A technical guide. Human-Wildlife Interactions Monograph, 1, 1–55.

71. Wickham H (2016) ggplot2: Elegant Graphics for Data Analysis. Springer-Verlag New York. ISBN 978-3-319-24277-4. Retrieved from https://ggplot2.tidyverse.org

72. Wilcox JT, Van Vuren DH (2009) Wild Pigs as Predators in Oak Woodlands of California. Journal of Mammalogy, 90(1), 114–118. 10.1644/08-mamm-a-017.1

73. Woodroffe R, Lindsey P, Romañach S, Stein A, Ole Ranah SM (2005) Livestock predation by endangered African wild dogs (Lycaonpictus) in northern Kenya. Biological Conservation, 124(2), 225–234. 10.1016/j.biocon.2005.01.028

