## Supplementary figures and images for "Converging paths: Harnessing ensemble modelling to predict human wild pig conflict risk zones in Tamil Nadu"

### The graph indicates the relative variable importance for correlation and AUC metrics.

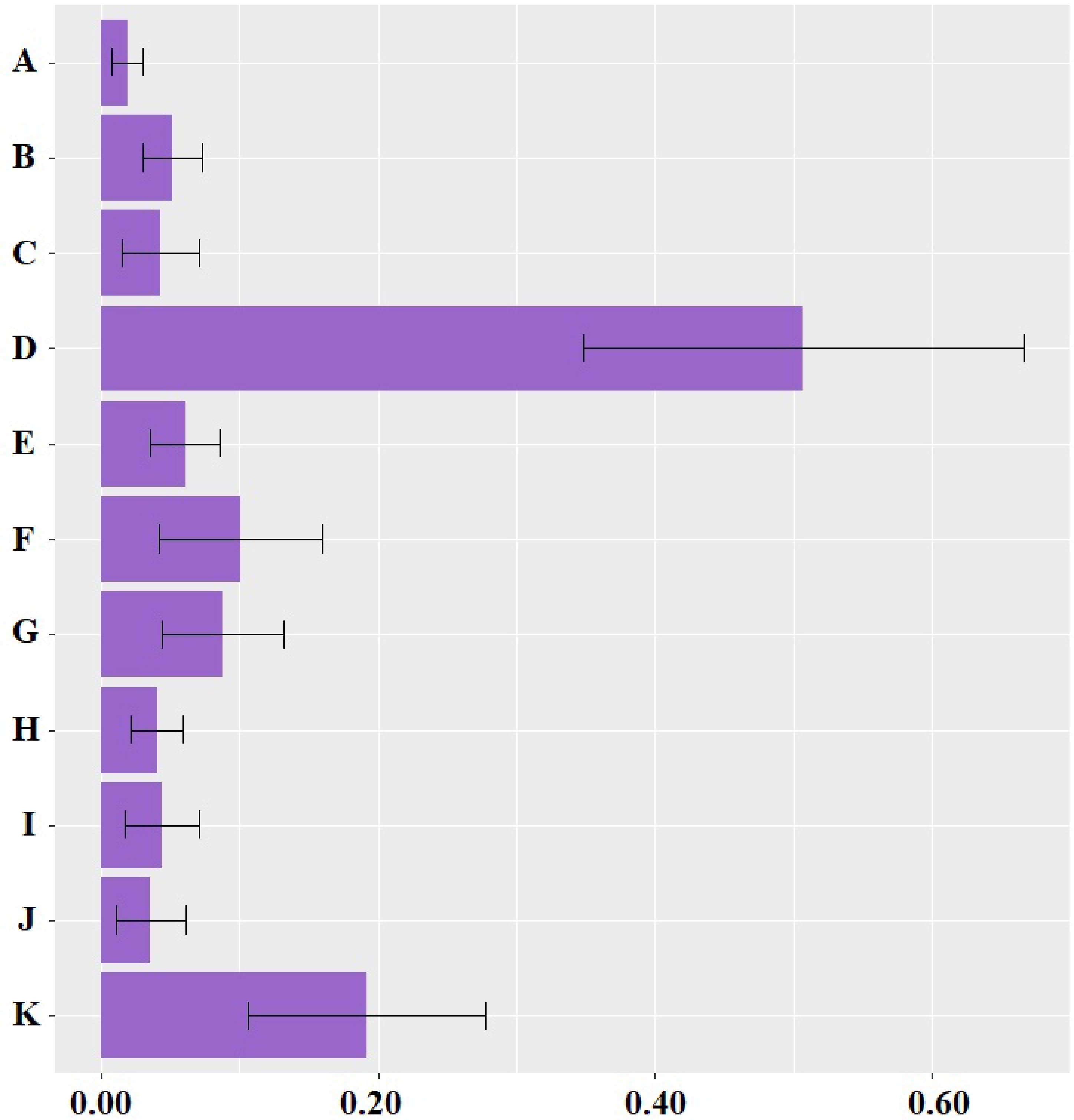

### The graph presents the Receiver Operating Characteristic (ROC) curve, depicting the performance of the model.

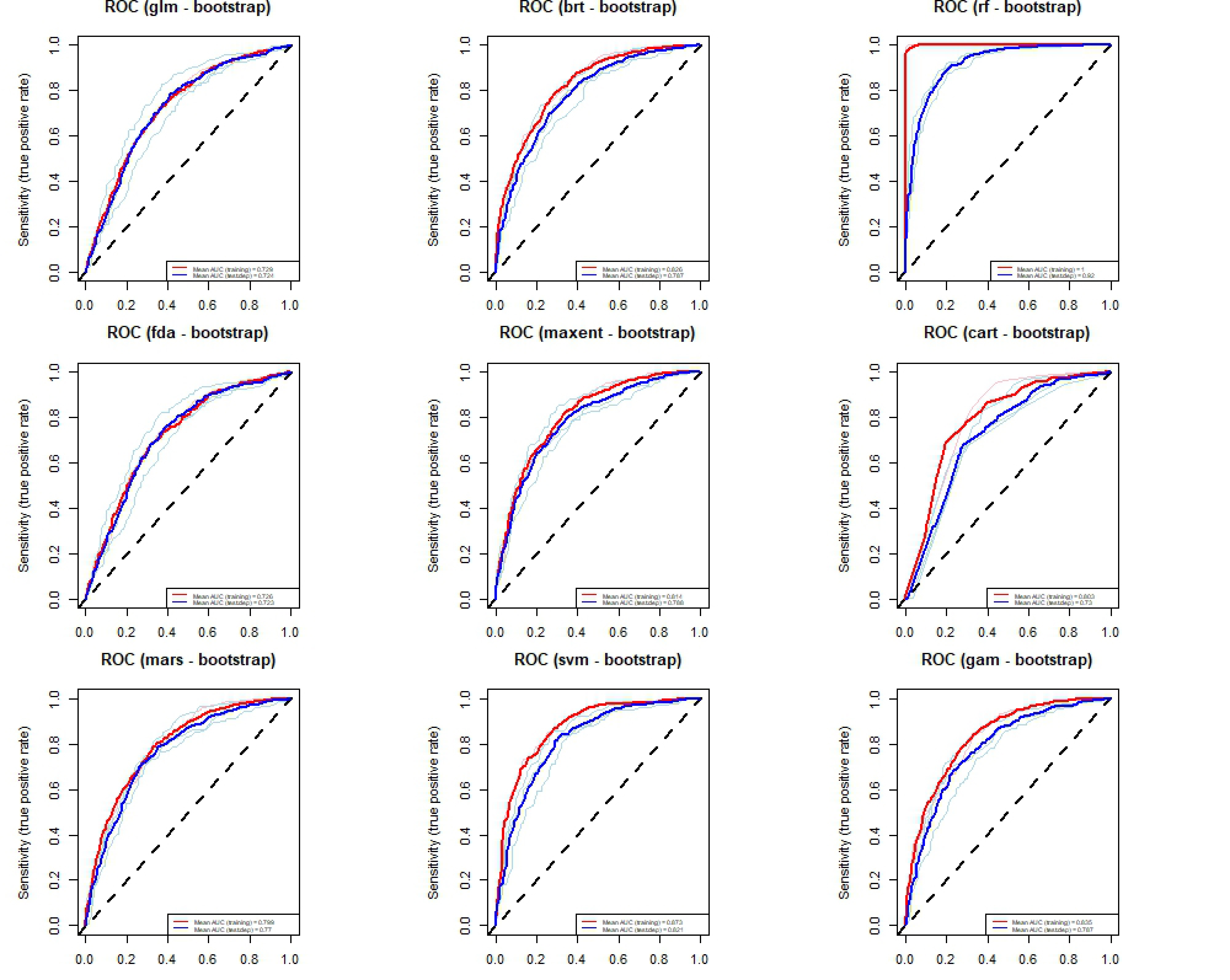

### The graph shows the Partial Response Curve, indicating the relationship between the independent variables and their impact on the prediction outcome.

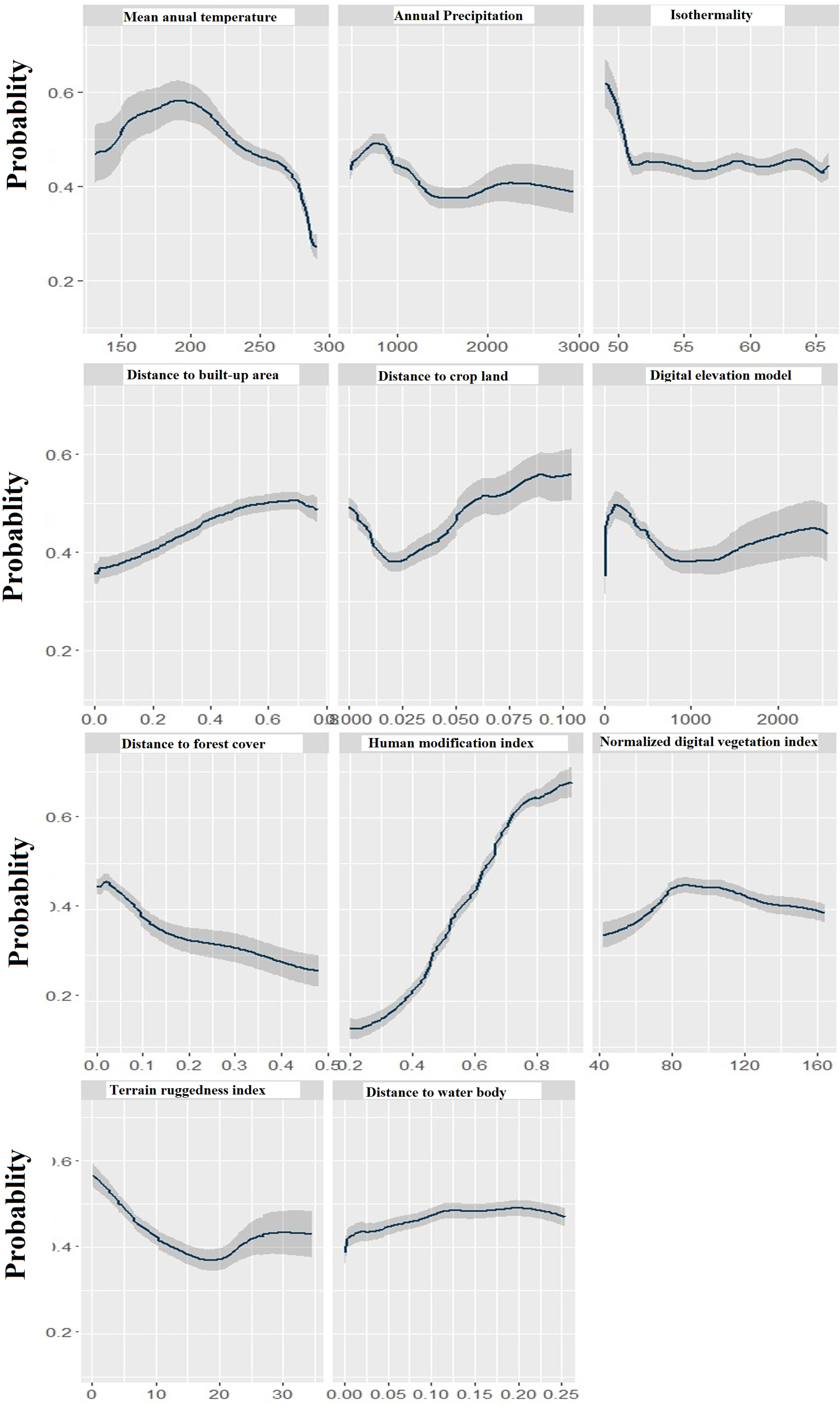
